# Trans-Radial Electrical Bioimpedance Velocimetry: A Novel Method for Detecting Cardiac Contractility

**DOI:** 10.1101/2022.06.05.494836

**Authors:** Alexandra I. Stump, Neil M. Dundon, Viktoriya Babenko, Alan Macy, Scott T. Grafton

## Abstract

Increasing insight into the complex human response to external states can be captured by measuring event-related cardiac sympathetic activity. However existing assays are either confounded by influence from other branches of the autonomic system, or require preprocessing steps that eliminate moment-to-moment capture of fluctuation. We accordingly tested a novel device (TREV) that measures cardiac impedance directly from the radial and ulnar arteries of the human forearm, while healthy human participants performed a small number of trials of a task known to elicit sympathetic drive, a maximum-strength grip task. TREV recorded robust estimates of contractility at each heartbeat, that allowed fully automated beatwise estimations. TREV further reliably described credible group-level departures from baseline aligned with each individual grip in the task. We conclude that the device can be a useful addition to a broadening field exploring event-related sympathetic perturbations.

## Introduction

An increasingly broad array of human sciences are gaining practical and theoretical value from noninvasively capturing event-related perturbations of the sympathetic branch of the autonomic nervous system. In earlier decades of research, electrocardiac assays of frequency-specific perturbations in heartrate were compromised by the confounding inputs of separate autonomic (e.g., parasympathetic) responses, even in lower frequency bands (Berntson et al., 1997; Reyes del Paso et al., 2013; Valenza et al., 2018). Later techniques used impedance assays via the thorax (impedance cardiogram (ICG)) and offered a more discrete capture of the sympathetic response, via estimates of cardiac contractility (Tomaka et al., 1997; Kelsey et al., 2004; Wright & Kirby, 2001). However such assays were first complicated by artefacts related to pulmonary activity and postural changes, and further restricted by difficulties identifying key events in the ICG, such as the opening of the aortic valve (a subtle perturbation (b-point); Cieslak et al., 2018). The latter impediments can be largely addressed using event-averaging, which indeed boosts signal-to-noise, but at the expense of sacrificing event-related precision.

More recent advancements in semi-automated preprocessing techniques of the ICG waverform have allowed increasingly event-related studies to emerge (Cieslak et al., 2018). Such studies have greatly advanced our understanding about how the sympathetic system is modulated when humans adapt to moment-to-moment fluctuations in their external state and enact processes such as motivation (Richter et al., 2016), cognitive effort (Kuipers et al., 2017) and optimal decision making (Dundon et al., 2020). In addition, event-related evidence further implicates the sympathetic system as an adaptive cortically-mediated allostatic response, and not simply reflexive homeostatic compensation shaped by brainstem reflex arcs (Dundon et al., 2020;2021, Babenko et al., 2022). Accordingly, additional assays of the sympathetic response continue to be explored, with specific focus on technologies that reduce preparation time, operate independently of the thorax and ultimately offer more robust contractility estimates at each heartbeat.

In the present study we therefore present event-related data recorded from a novel thorax-independent device that measures cardiac impedance directly from the radial and ulnar arteries of the human forearm (Macy and Bernstein, 2020). Using an easily applied set of electrode wristbands (four electrodes in total, see Figure 1A), the device detects changes in resistance that occur as intraluminal red blood cells align in response to a pressure wave transmitted through the arterial tree at the opening of the aortic valve, prior to bulk blood flow and its associated pulse wave. The pressure wave is directly ascribed to cardiac contractional vigor. The cellular alignment causes a drop in impedance which is monitored continuously by the device’s electrodes. The third derivative (“jerk”) of the continuously recorded impendence can therefore offer a robust estimate of contractility, i.e., how fast impedance acceleration changes with respect to time (Bernstein et al., 2015).

**Figure 1.**
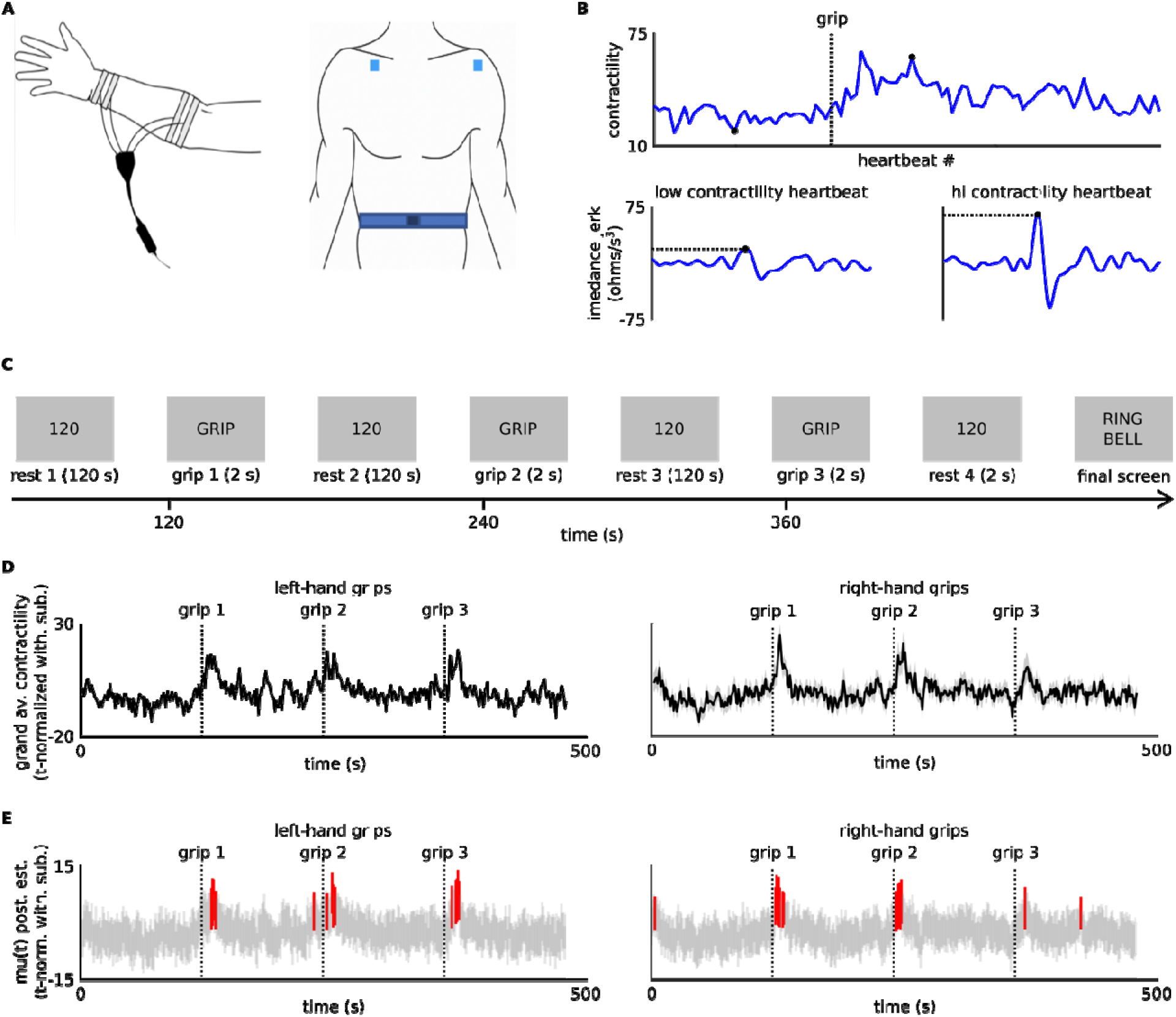
TREV reliably identifies contractility departing baseline while partipants make grips. **A.** Electrode placement for TREV (left) and EKG (right). **B.** Exemplar contractility timeseries (top) of the 200^th^ to 300^th^ heartbeat for a single subject during a grip with their right hand. Contractility epochs (bottom) for a heartbeat with low and hi contractility. Each epoch spans 700 ms relative to the EKG R-point and depicts the third derivative of impedance (jerk) recorded with TREV. Note the differences in max amplitudes between epochs. **C.** Task outline. Participants completed three incentivized grips with each of their right and left hand, interlaced with two-minute rests. **D.** Grand averaged contractility across participants. Contractility was first t-normalized within-subjects and separately for left and right-hand grips. Shaded grey areas depict the standard error of the mean across subjects at each timepoint. **E.** Posterior estimates of group-mean contractility at each timepoint (mu(t)). Vertical grey lines depict the highest density interval (HDI) at each time point and where this HDI is credibly above the mean across all timepoints (M_mu_: see methods) the line is red. Note the different scales between panels D and E, as the hierarchal model (panel E) constrains mu(t) estimates, making it inherently conservative (see methods). Dashed vertical lines in panels D and E reflect the cue to grip in each case.

## Materials and Methods

### Participants and experimental overview

Thirty one healthy humans (19 females, 12 males) participated in the study after providing informed consent in accordance with the University of California, Santa Barbara (UCSB) Institutional Review Board. Participants self-reported no cardiovascular abnormalities. The average age of participants was 23.4 years. One participant was excluded due to excessively noisy data, leaving a final sample of n=30. Participants were compensated $10/hour plus a potential additional $10 bonus depending on task performance (see Grip Task below).

Participants performed two blocks of a maximum grip task (Grip Task), while three simultaneous physiology timeseries were recorded. The first timeseries was time-varying cardiac impedance derived from trans-radial electrical bioimpedance velocimetry (TREV; Macy & Bernstein, 2020) electrodes attached to the forearm contralateral to the hand administering grips. The second timeseries was a standard electrocardiogram (EKG). The last timeseries recorded the continuous respiration cycle.

### Apparatus

TREV electrodes, were amplified by an NICO100D (BIOPAC) smart amplifier. Electrocardiogram electrodes were amplified by an EKG100D (BIOPAC) smart amplifier. Respiration cycle was recorded using a TSD221 (BIOPAC) respiration belt. Force exerted in the Grip Task was recorded using a SS56L (BIOPAC) grip bulb. All continuous signals were integrated using an MP160 (BIOPAC) amplifier and processed online using BIOPAC AcqKnowledge software (BIOPAC). Visual stimuli were presented using Microsoft Powerpoint. Offline preprocessing of recorded timeseries was conducted using the Moving Ensemble-Average Pipeline (MEAP; Cieslak et al., 2018) and MATLAB. Bayes models were fitted using No U-Turn sampling (NUTS) Hamiltonian Monte Carlo, fitted with PyMC3 Python3 functions (Salvatier et al., 2016).

### General Procedure

All data were recorded in a single session lasting approximately 45 minutes (including initial equipment setup). Participants first washed their hands and forearms with water and regular soap to remove dirt or oily residues. In a private setup room, an experimenter then placed four TREV electrodes on the forearm contralateral to the grip hand of the first block (see Grip Task, below) as depicted in Figure 1A: two ventrally on the distal part of the forearm, just below where the wrist meets the hand, and two on the proximal part of the forearm, just below where the elbow meets the forearm. Each electrode pair was spaced one centimeter apart. The experimenter then placed two electrocardiogram electrodes on the participant’s chest: one below the right collarbone and one where the deltoid meets the chest (see Figure 1A). The experimenter then brought participants to the testing room, attached electrodes to their associated amplifiers, attached the respiration belt to their abdomen and seated them at the testing table approximately one meter from a computer screen. Once seated in the testing room, participants learned how to properly hold and squeeze the grip bulb, with the tubing facing down and in a manner that involved the whole hand. The experimenter then asked participants to grip the bulb as hard as possible with each hand, recording each value (max thresholds).

### Grip Task

We administered the Grip Task to capitalize on the known increase in sympathetic activation while humans apply maximum grip force (Richter, 2015; Richter, Gendolla & Wright, 2016; Stanek & Richter, 2015, 2016, 2021). Participants performed two blocks of three trials, gripping with a different hand in each block (block-hand order was determined with uniform (p=0.50) probability for each participant). After recording participants’ maximum forces (max thresholds; above), the experimenter then explained the experimental protocol, which is depicted in Figure 1C. Trials began with an on-screen countdown timer, which counted down through a two-minute rest period to a “go” cue. Once the go cue appeared, participants would squeeze the bulb as hard as possible for two seconds. The countdown period of the next trial’s rest period then immediately began. Once the experimenter started recording the physiology data, they loaded the visual stimuli and left the room. Before starting each block, participants sat still for three minutes. At the end of the third trial on each block, a timer counted down to a visual stimulus that instructed participants to ring a bell to alert the experimenter they had finished. Each of the three trials was therefore preceded and followed by a two-minute rest. To incentivize participants to grip with maximum strength, we imposed a bonus system, whereby participants who reached a threshold of ± 0.04 Kgf/m^2^ of their hand-specific max-thresholds on any grips would win a $10 bonus. The experimenter disclosed this rule to participants after recording the max thresholds and did not inform participants if they had achieved the bonus until after all testing was completed. After completing the first block, the experimenter transferred the TREV electrodes to the other arm.

### Cardiovascular preprocessing

During recording, the AcqKnowledge software applied an online lowpass filter (max cutoff = 20 Hz) to raw cardiac impedance timeseries recorded by the TREV electrodes, and then took its 3^rd^ derivative (i.e., “Jerk”) as a continuous estimation of contractility (Bernstein et al., 2015). This raw contractility timeseries was then imported together with the raw EKG and respiration timeseries to the MEAP software for minimal offline processing. MEAP first automatically labelled the R-peaks of the EKG timeseries, which we used as an index for the moment in time of each individual heartbeat. We next used these R-peak time indices to extract epochs spanning 700 ms around each heartbeat from the raw contractility timeseries (contractility epochs). MEAP also computed estimates of heart-rate at each beat from the R-peaks. MEAP outputs were then transferred to MATLAB, where the maximum amplitude in each contractility epoch was computed as an estimation of each heartbeat’s contractility (beatwise contractility timeseries). Then, separately for each subject, and each block, we conducted an additional regression procedure (Dundon et al., 2020) to remove the additional confounding effects of heart-rate and respiration from the beatwise contractility timeseries. In a multiple regression model, we regressed the vector beatwise contractility as a function of an intercept and three regressors: (i) the phase of respiration at each heartbeat, (ii) the amplitude of respiration at each heartbeat and (iii) the heartrate at each heartbeat. To down sample each regressor to beatwise estimates, we used the value from raw timeseries closest to the time of each R-peaks. We added the estimated intercept to the residuals from this model as the “residualized” contractility timeseries, i.e., with the effects of the above three regressors removed. Given both between-subject and within-subject variation in heart rate, we next applied temporal resampling of each block’s residualized timeseries to allow meaningful comparisons across participants. For this, we used onedimensional linear interpolation across time to recreate residualized timeseries sampled at equal time intervals. Specifically, we took 479 estimates, spaced exactly one second apart, from 2 seconds post block onset until 480 seconds post block onset (interpolated contractility timeseries). Finally, these interpolated contractility timeseries were normalized as a t-statistic, i.e., each interpolated contractility estimate expressed as a t-statistic relative to the timeseries’s remaining 478 values. We will refer to this t-statistic-normalized timeseries from now on as the “contractility” timeseries. A grand average contractility timeseries across participants, separately for each block, is presented in Figure 1D.

### Bayesian modeling framework

The primary objective of our analyses was to determine whether TREV could reliably capture increases in group-level contractility that corresponded to the events in the grip task. For this, we used a hierarchical Bayesian framework which hypothesized that the (n=30) group distribution of contractility estimates at each timepoint (t) formed a Student’s T distribution, T(t)~Student’s T(mu(t),sig(t),nu). We formally considered contractility to have increased beyond baseline at a given moment where the estimated mean of a timepoint’s distribution (mu(t)) credibly exceeded the mean across all timepoints (M_mu_). M_mu_ is itself fitted in the same model as the mean of a hierarchical Gaussian distribution (G_mu_) which constrains estimates of each mu(t) by serving as their prior (G_mu_~N(M_mu_,S_mu_)). Given how Bayes theorem ascribes joint probabilities to both the prior and the observed data in posterior estimates, this distributional hierarchical framework is inherently conservative with respect to type one error for each estimate of mu(t). For example, if most values for mu(t) are within a tight range (as we would expect in a dataset of contractility values with long rest periods between grips), the hierarchical distribution will be characterized by a more certain mean and low variance (low value of S_mu_), which would then serve as a strict prior on mu(t) estimates, biasing them toward the group mean (i.e., a nail that stands out gets hammered in). This hierarchical framework therefore requires strong evidence before any mu(t) is formally accepted as a credible departure. In other words, in a context requiring multiple hypothesis tests, the hierarchical Bayesian framework imposes an adjustment to the level of evidence needed for credible effects, where the data itself determines that level of adjustment instead of an arbitrary criterion (e.g., Bonferroni).

We fitted a hierarchical model separately for blocks where grip was administered with the right and left hand. In each case, the specific free parameters of our model were: mu(t) and sigma(t), i.e., the 479 timepoint-specific mean and standard deviation parameters for group-level SNS distributions at each timepoint across each block. We did not fit the nu parameter hierarchically and assigned it the same uninformed prior (nu=1) in each model. As mentioned above, each mu(t) parameter was constrained by a hierarchical Gaussian distribution (G_mu_) with free parameters M_mu_ and S_mu_ corresponding respectively to its mean and standard deviation. M_mu_ was assigned an uninformed Gaussian prior, N(0,1), while S_mu_ was assigned an uninformed half-Gaussian prior (forcing values to be positive), halfN(1). Each sigma(t) was also constrained by hierarchical Gaussian distribution (G_sigma_), which respectively used an uninformed Gaussian and half-Gaussian prior for its two free parameters, i.e., its mean (M_sigma_~N(0,1)) and standard deviation (S_sigma_~halfN(1)). We formally compared each mu(t) posterior with that of the M_mu_ by computing the minimum-width Bayesian credible interval (Highest Density Interval (HDI)) of mu(t) - M_mu_ and only considered strong evidence of a departure at each timepoint, i.e., where resulting HDIs did not contain zero.

Contractility timeseries were z-score normalized prior to fitting across all participants. Each model’s posterior distributions were sampled across four chains of 5000 samples (20000 total), after burning an initial 5000 samples per chain to tune the sampler’s step-size to reach 0.95 acceptance. We estimated HDIs using the default setting in the arviz package (Kumar et al., 2019).

## Results

We tested whether a thorax-independent monitor of cardiac impedance (TREV) could reliably describe fluctuations in cardiac contractility that credibly exceed baseline as human participants perform a task known to drive increased cardiovascular sympathetic stress. 30 participants completed two blocks of three incentivized max-intensity grips, with rest periods of two minutes both between each grip and following the final grip. We simultaneously recorded continuous cardiac impedance from the radial and ulnar artery contralateral to the gripping hand using TREV, and following minimal preprocessing, computed beatwise measures of contractility as the third derivative of this cardiac impedance (“jerk”).

Participants understood and performed the behavioral requirements of the task. Participants also showed strong motivation to grip at maximum intensity, supported by 29 out of 30 achieving a bonus payment (contingent on beating their predetermined max threshold) with at least one hand, and 21 out of 30 achieving the bonus payment with both.

Figure 1B depicts how the contractility at each heartbeat is derived from the maximum impedance jerk across its epoch (see contractility epoch, in methods). Two exemplar heartbeats are shown from a single subject, one in the rest phase prior to the second grip with their right hand and another in the grip’s immediate aftermath. In contrast to labeling b-points in ICG preprocessing, the robust jerk measures allowed maximum values to be estimated in an automated pipeline with no human involvement.

We next used linear resampling to temporally align contractility across participants, and normalized each block separately as a t-statistic. Grand-averages of these resampled, normalized contractility waveforms across subjects are depicted in Figures 1D. Visual inspection of these grand averages show marked increase in group-level contractility in close temporal approximation to each grip.

The results of the hierarchical Bayesian model fitted to contractility timeseries accompanying left-hand grips are depicted in the left panel of Figure 1E. TREV reliably captured contractility exceeding baseline following grips with the left hand. Left hand grips were accompanied by credible baseline departure at timepoints [131, 132, 133, 135, 232, 245, 250, 251, 253, 368, 372, 373, 374, 375], in seconds relative to the onset of the first trial. Expressing these timepoints in seconds relative to the nearest grip, they occurred at grip 1: [11, 12, 13, 15], grip 2: [-8, 5, 10, 11, 13] and grip 3: [8, 12, 13, 14, 15]. Each grip was therefore accompanied by at least 4 individual seconds of credible baseline departure. Departures mostly followed the grips and never followed a grip by more than 15 seconds. Each grip was associated with at least two consecutive seconds of baseline departure, with grip 3 associated with the longest sustained peak contractility (four consecutive points).

The results of the hierarchical Bayesian model fitted to contractility timeseries accompanying right-hand grips are depicted in the right panel of Figure 1E. Right-hand grips were accompanied by credible baseline departure at timepoints [6, 125, 126, 127, 128, 129, 132, 133, 244, 245, 246, 247, 248, 249, 371, 426], in seconds relative to the onset of the first trial. Expressing these timepoints in seconds relative to the nearest grip, they occurred at grip 1: [-114, 5, 6, 7, 8, 9, 12, 13], grip 2: [4, 5, 6, 7, 8, 9] and grip 3: [11, 66]. Discounting the two outliers (preceding grip 1 and following grip 3), each grip was therefore accompanied by at least 1 second of credible baseline departure. Departures all followed the grips and never followed a grip by more than 13 seconds. Grip 2 was associated with the longest sustained peak contractility (six consecutive points). TREV again appeared to reliably capture contractility exceeding baseline following grips with the right hand, although a pair of outliers were present and the duration of peak contractility seemed to abate over the course of the three grips.

## Discussion

There is expanding interest across multiple human research disciplines in robustly capturing event-related perturbations of the sympathetic stress response. With this there a need for new assays of cardiac contractility that both reduce preparatory requirements and offer increased signal strength in the face of background noise. In this study we accordingly used a novel transradial electrical bioimpedance velocimetry device (TREV), attached to the forearm of human participants, and investigated whether it could reliably capture changes in group-level contractility that corresponded to the events known to increase sympathetic drive, a max grip task (Grip Task). We observed that TREV electrodes can be applied in a matter of minutes with minimal training and preparation, and can even be repositioned (from one arm to the other) efficiently between blocks of a task. We further observed TREV to register easily identifiable (by eye) beat-to-beat signals from the radial and ulnar artery corresponding to the third derivative of cardiac impedance (“jerk”). In preprocessing, we controlled for the confounding effects of respiratory activity and heart rate on beatwise contractility timeseries. Then, using a hierarchical Bayesian framework, we observed these contractility timeseries to reliably depart baseline at key events in the Grip Task. Remarkably, these departures were seen at the single-trial level across participants (i.e., without averaging across trials). We therefore conclude that TREV offers an exciting development in cardiac autonomic stress research for human researchers interested in event-related capture of cardiac contractility.

We employed a data-driven analysis framework, which accommodated the entire timeseries of data recorded across sessions, to determine when contractility estimated by TREV credibly exceeded baseline fluctuations. The primary advantage of this framework is that it removed all need to impose arbitrary criteria on grip events or contractility activity, i.e., a priori deciding epochs around task events to refine analysis, or a priori deciding a criterion that constituted “credibly exceeding baseline”. The analysis was further not assisted by any averaging across events to reduce signal-to-noise. The hierarchical Bayesian framework also imposed conservativeness with respect to credible departures from baseline across a large number of hypothesis tests. We nonetheless revealed reliable group-level increases in contractility at each of the six grips executed by participants. Note that our criterion for baseline was the average value across all datapoints in the timeseries, which theoretically incorporates all preparatory increases sympathetic activity leading up to grip execution. Thus, while our framework focused on identifying robust measures of peak contractility, an exciting avenue for future studies might be to implement machine learning techniques on TREV data recorded simultaneous to tasks such as the Grip Task to probe the device’s ability to track when humans begin ramping up preparatory stress responses to effortful action, and not just identify moments of departure from baseline.

We additionally observed differences between the contractility profiles elicited by grips with the left and right hand. For grips with the left hand, contractility only exceeded baseline within a 15-second window on either side of grip execution, and exceeded baseline for a similar number of timepoints (four or five) across grips. In contrast, a pair of outliers emerged in the contractility timeseries accompanying right-hand grips, and further, the number of timepoints exceeding baseline appeared to diminish across grips. Taking the latter observation first, this reduction in baseline departures across right-hand grips could be a result of participants adapting to the known effort requirements of grips with their more dominant hand. In terms of timing, it is interesting to note that for almost the entirety of the two minute preparatory warning period, there was no evidence for anticipatory changes of contractility until just prior to commencement of the grip. That is, the cortical regulation of sympathetic drive in this task may be adopting a learned strategy of waiting until the last moment to enhance cardiac inotropy.

Motivational intensity is known to scale with the level of task difficulty, an effect which has been observed in both cognitive and grip tasks (see: Richter et al., 2016 for a review). Participants may therefore require less intense mobilization for later dominant grips, as they amass more accurate feedforward control due to more efficiently formed internal models of task-appropriate limb dynamics (Heuer, 2006). In a similar vein, participants may also have been more confident that they had achieved the bonus in the first two dominant grips (i.e., more so than in the non-dominant grips), lowering the incentive to grip more intensely in the third dominant grip. In terms of the outliers, the departure from baseline at the very beginning of the timeseries of right-hand grips is consistent with SNS activity increasing early in tasks while humans adjust to situational requirements (Dundon et al., 2020), and this again might be more relevant for the dominant hand.

In conclusion, we here observe that thorax-independent TREV reliably captured contractility increases to individual events, and offers considerable advantages for capturing event-related cardiac responses in more generalized real-world task settings. Such capture of contractility signals would greatly contribute toward improving our knowledge of how humans synchronize sympathetic state while monitoring broader state information, allowing us to develop more holistic technologies for human-machine integration that can assist with situational awareness, maneuverability and decision making.

## Acknowledgements

Supported by the Institute for Collaborative Biotechnologies under Cooperative Agreement W911NF⍰19⍰2⍰0026 and grant W911NF⍰16⍰1⍰0474, both from the Army Research Office.

## References

Babenko, V., Dundon, N. M., Stump, A. I., Turbow, M., Cieslak, M., Macy, A., & Grafton, S. T. (2022). An accelerometer based heart monitor to measure changes of the autonomic nervous system. BioRxiv. https://doi.org/10.1101/2022.06.02.494453

Bernstein, D. P., Henry, I. C., Lemmens, H. J., Chaltas, J. L., DeMaria, A. N., Moon, J. B., & Kahn, A. M. (2015). Validation of stroke volume and cardiac output by electrical interrogation of the brachial artery in normals: assessment of strengths, limitations, and sources of error. Journal of Clinical Monitoring and Computing, 29(6), 789–800.

Berntson, G. G., Thomas Bigger Jr, J., Eckberg, D. L., Grossman, P., Kaufmann, P. G., Malik, M., … & Van Der Molen, M. W. (1997). Heart rate variability: origins, methods, and interpretive caveats. Psychophysiology, 34(6), 623–648.

BIOPAC Systems, Inc, Goleta, CA

Dundon, N. M., Garrett, N., Babenko, V., Cieslak, M., Daw, N. D., & Grafton, S. T. (2020). Sympathetic involvement in time-constrained sequential foraging. Cognitive, Affective, & Behavioral Neuroscience, 20(4), 730–745.

Dundon, N. M., Shapiro, A. D., Babenko, V., Okafor, G. N., & Grafton, S. T. (2021). Ventromedial prefrontal cortex activity and sympathetic allostasis during value-based ambivalence. Frontiers in behavioral neuroscience, 15.

Cieslak, M., Ryan, W. S., Babenko, V., Erro, H., Rathbun, Z. M., Meiring, W., … & Grafton, S. T. (2018). Quantifying rapid changes in cardiovascular state with a moving ensemble average. Psychophysiology, 55(4).

Heuer, H., 2007. Control of the dominant and nondominant hand: exploitation and taming of nonmuscular forces. Experimental brain research, 178(3), pp.363–373.

Kelsey, R. M., Soderlund, K., & Arthur, C. M. (2004). Cardiovascular reactivity and adaptation to recurrent psychological stress: Replication and extension. Psychophysiology, 41(6), 924–934.

Kumar, R., Carroll, C., Hartikainen, A. & Martin, O. (2019). ArviZ a unified library for exploratory analysis of Bayesian models in Python. J. Open Source Softw. 4, 1143.

Macy, A., & Bernstein, D. (2020). Trans-radial electrical bioimpedance velocimetry (TREV). BIOPAC.

Reyes del Paso, G. A., Langewitz, W., Mulder, L. J., Van Roon, A., & Duschek, S. (2013). The utility of low frequency heart rate variability as an index of sympathetic cardiac tone: a review with emphasis on a reanalysis of previous studies. Psychophysiology, 50(5), 477–487.

Richter, M. (2015). Goal pursuit and energy conservation: energy investment increases with task demand but does not equal it. Motivation and Emotion, 39, 25–33.

Richter, M., Gendolla, G., & Wright R. (2016). Chapter Five - Three Decades of Research on Motivational Intensity Theory: What We Have Learned About Effort and What We Still Don’t Know. Elselvier, 3, 149–186. https://doi.org/10.1016/bs.adms.2016.02.001

Salvatier, J., Wiecki, T. V., & Fonnesbeck, C. (2016). Probabilistic programming in Python using PyMC3. PeerJ Computer Science, 2, e55.

Stanek, J. C., & Richter, M. (2015). Energy investment and motivation: the joint impact of task demand and reward value on exerted force in hand grip tasks (in press).

Stanek, J. C., & Richter, M. (2016). Evidence against the primacy of energy conservation: Exerted force in possible and impossible handgrip tasks. Motivational Science, 2, 49–65.

Stanek, J. C., & Richter, M. (2021). Energy investment and motivation: The additive impact of task demand and reward value on exerted force in hand grip tasks. Motivation and Emotion, 45, 131–145.

Tomaka, J., Blascovich, J., Kibler, J., & Ernst, J. M. (1997). Cognitive and physiological antecedents of threat and challenge appraisal. Journal of Personality and Social Psychology, 73(1), 63.

Valenza, G., Citi, L., Saul, J. P., & Barbieri, R. (2018). Measures of sympathetic and parasympathetic autonomic outflow from heartbeat dynamics. Journal of Applied Physiology, 125(1), 19–39.

Wright, R. A., & Kirby, L. D. (2001). Effort determination of cardiovascular response: An integrative analysis with applications in social psychology. Advances in Experimental Social Psychology, 33, 255–307.

